# Diversity of Environmental and Genome-Associated Eukaryotic Circular Rep-Encoding Single-Stranded DNA Viruses

**DOI:** 10.64898/2026.11.20.768052

**Authors:** Oliver K. Bezuidt, Frederik Schulz

**Affiliations:** Department of Biochemistry, Genetics and Microbiology, University of Pretoria, Pretoria, South Africa; DOE Joint Genome Institute, Lawrence Berkeley National Laboratory, Berkeley, CA, 94720, USA

## Abstract

Circular Rep-encoding single-stranded (CRESS) DNA viruses are widespread across diverse environments, yet their genomic diversity, evolutionary relationships, and associations with eukaryotic hosts remain poorly understood. Here, we investigated the evolutionary diversity of CRESS DNA viruses by integrating replication-associated protein (Rep) sequences from environmental viral metagenomes, publicly available viral references, and endogenous viral elements (EVEs) identified in fungal and algal genomes. Genome-wide screening revealed numerous EVEs distributed across diverse fungal and algal lineages, highlighting the potential historical associations between CRESS DNA viruses and eukaryotic hosts. Phylogenetic analyses revealed considerable diversity among CRESS DNA viruses, including sequences that could not be assigned to currently recognized viral families. Yet, several of these unclassified sequences formed well-supported monophyletic clades, suggesting that existing taxonomic classifications may require further revision. Furthermore, phylogenetic relationships within and among established viral families suggest that existing taxonomic boundaries may not fully capture the evolutionary diversity of these viruses. The phylogenetic placement of fungal and algal EVEs alongside environmental and reference viral sequences further provides insights into the evolutionary relationships between CRESS DNA viruses and eukaryotic genomes. Collectively, our findings expand the known evolutionary diversity of CRESS DNA viruses, identify distinct viral lineages requiring further taxonomic evaluation, and provide a framework for investigating their genomic evolution, taxonomic relationships, and associations with eukaryotic hosts.

## Introduction

Circular single-stranded DNA (ssDNA) viruses that encode a replication-associated protein (Rep) include some of the smallest autonomously replicating viral genomes. These viruses belong to the phylum Cressdnaviricota and are commonly referred to as circular Rep-encoding single-stranded (CRESS) DNA viruses (Rosario et al., 2012, Krupovic et al., 2020). Like many viral groups uncovered through environmental sequencing, CRESS DNA viruses are exceptionally diverse and infect a broad spectrum of eukaryotic hosts, including plants, animals, and fungi (Rosario et al., 2017, Zhao et al., 2019). They have also been detected in a wide array of environmental samples, indicating a global distribution across terrestrial, freshwater, and other natural ecosystems (Simmonds et al., 2017, Kazlauskas et al., 2019)

Despite their ubiquity, the diversity, host range, and ecological roles of CRESS DNA viruses remain poorly understood. Historically, virology has focused on viruses that cause disease or have economic importance, resulting in greater attention being given to pathogenic ssDNA viruses, such as those in the *Geminiviridae* and *Circoviridae* families. These families comprise several agriculturally important plant and animal pathogens. In contrast, environmentally derived or poorly characterized host-associated CRESS viruses remain largely unexplored. Small genomes, high sequence divergence, and sparse representation in reference databases limit the detection of CRESS DNA viruses by conventional virological and sequence-based methods (Krishnamurthy and Wang, 2017). At the same time, large-scale metagenomic sequencing efforts have revealed that the number of viral sequences recovered from environmental samples far exceeds the number of experimentally characterized virus isolates. This has uncovered a vast “dark matter” of viral diversity that traditional tools have overlooked (Santiago-Rodriguez and Hollister, 2022, Roux and Coclet, 2026).

The advent of next-generation sequencing (NGS) and metagenomic approaches has begun to transform our understanding of viral diversity, including that of CRESS DNA viruses. High-throughput metagenomic sequencing now routinely uncovers novel circoviruses, circovirus-like genomes, and other CRESS DNA viruses from a wide range of habitats, including soils, freshwater systems, wastewater, and host-associated microbiomes (Delwart and Li, 2012, Rosario et al., 2017, Zhao et al., 2019). Many of these viruses would likely have remained undetected using classical methods. These discoveries have revealed extensive genomic diversity, identified previously unrecognized host associations, and raised important questions about viral origins, genome organization, and ecological roles. Continued improvements in sequencing methods, data analysis, and quality control will be essential for studying CRESS DNA viruses more accurately. These advances will help clarify their environmental distribution, host interactions, and ecological roles. Previous studies have shown that CRESS DNA viruses may occasionally become integrated into host genomes, leaving genetic signatures known as endogenous viral elements (EVEs). Although the mechanisms by which non-retroviral viruses, including members of the Cressdnaviricota, integrate remain unclear (Krupovic and Forterre, 2015), expanded eukaryotic genome sequencing has increased the detection of EVEs across diverse eukaryotic taxa (Aswad and Katzourakis, 2012, Krupovic et al., 2020). Endogenous sequences related to circoviruses and other CRESS DNA viruses have been reported in animal genomes, consistent with long-term virus-host associations (Dennis et al., 2019, Katzourakis and Gifford, 2010). Validated EVEs can identify historical virus-host associations that are absent from contemporary sampling (Dennis et al., 2018). Together, these endogenous elements provide insights into viral evolution and long-term virus–host interactions, while complementing environmental metagenomic studies by linking current viral diversity to past viral infections.

Collectively, environmental metagenomics and EVE-based analyses provide complementary perspectives for reconstructing the ecological and evolutionary trajectories of CRESS DNA viruses, offering a more complete picture of their diversity, host interactions, and long-term evolutionary dynamics (Holmes, 2011, Aswad and Katzourakis, 2012, Rosario et al., 2017). In this study, we characterize CRESS DNA virus-related Rep sequence diversity in IMG/VR and screen fungal and protist (including algal) genome assemblies from MycoCosm and PhycoCosm databases for genome-associated Rep-like loci. We place environmental, genome-associated, and ICTV reference viral sequences in a common phylogeny to examine sequence relationships and provisional family associations. By combining contemporary environmental viral sequences with endogenous viral records preserved within eukaryotic genomes, this study aims to improve our understanding of the diversity, host associations, and evolutionary relationships of CRESS DNA viruses across environmental and eukaryotic systems.

## Materials and Methods

### Sequence data acquisition and processing

#### Acquisition and processing of Rep protein sequences

A total of 61,709 protein sequences associated with the phylum Cressdnaviricota were retrieved from UniProt (accessed September 2024). We filtered these to 11,892 complete sequences annotated as replication-associated protein (Rep) and containing the conserved rolling-circle replication motifs RCR-1, RCR-2, and RCR-3. These were clustered into 3,115 Rep representative sequences with cd-hit v.4.8.1 (-c 0.95 and -aS 0.8). To recruit additional homologs, we obtained 108,138 Cressdnaviricota protein sequences from GenBank (accessed September 2024). These were queried by the 3,115 representative Rep proteins using jackhmmer v3.4 in 5 iterations (-E 1e-10 -N 5). The search resulted in a set of 38,191 Rep proteins which were also clustered into 4,467 complete proteins with cd-hit v.4.8.1 (-c 0.95 and -aS 0.8) and merged with the initial 3,115 Rep proteins from UniProt, resulting in a total of 7,582 representative sequences.

To identify additional Rep proteins, we queried the merged sequences against IMG/VR viral datasets (IMGVR_all_proteins.faa: 220,779,163 protein sequences and IMGVR_all_proteins-high_confidence.faa: 112,567,455 protein sequences, accessed October 2024) using jackhmmer (-E 1e-10 -N 5). After dereplication (cd-hit -c 0.95 and -aS 0.8), searches of the IMGVR_all_proteins_high_confidence.faa and IMGVR_all_proteins.faa data yielded 11,819 and 23,689 Rep proteins, respectively. We merged and dereplicated both hit sets, yielding 25,416 candidate sequences for validation with HHsearch v.3.3.0 against the 2022 PDB70 database (-E 1e-5). We retained sequences as high-confidence Rep proteins when HHsearch returned a match with probability >= 90% to an entry annotated as a Rolling circle replication, Rep protein/domain, replicase-associated protein, or Replication associated protein (hereafter referred to as high-confidence Rep proteins). The retained high-confidence Rep proteins were 25,416 in total (see **Figure 1** for the pipeline workflow).

**Figure 1:**
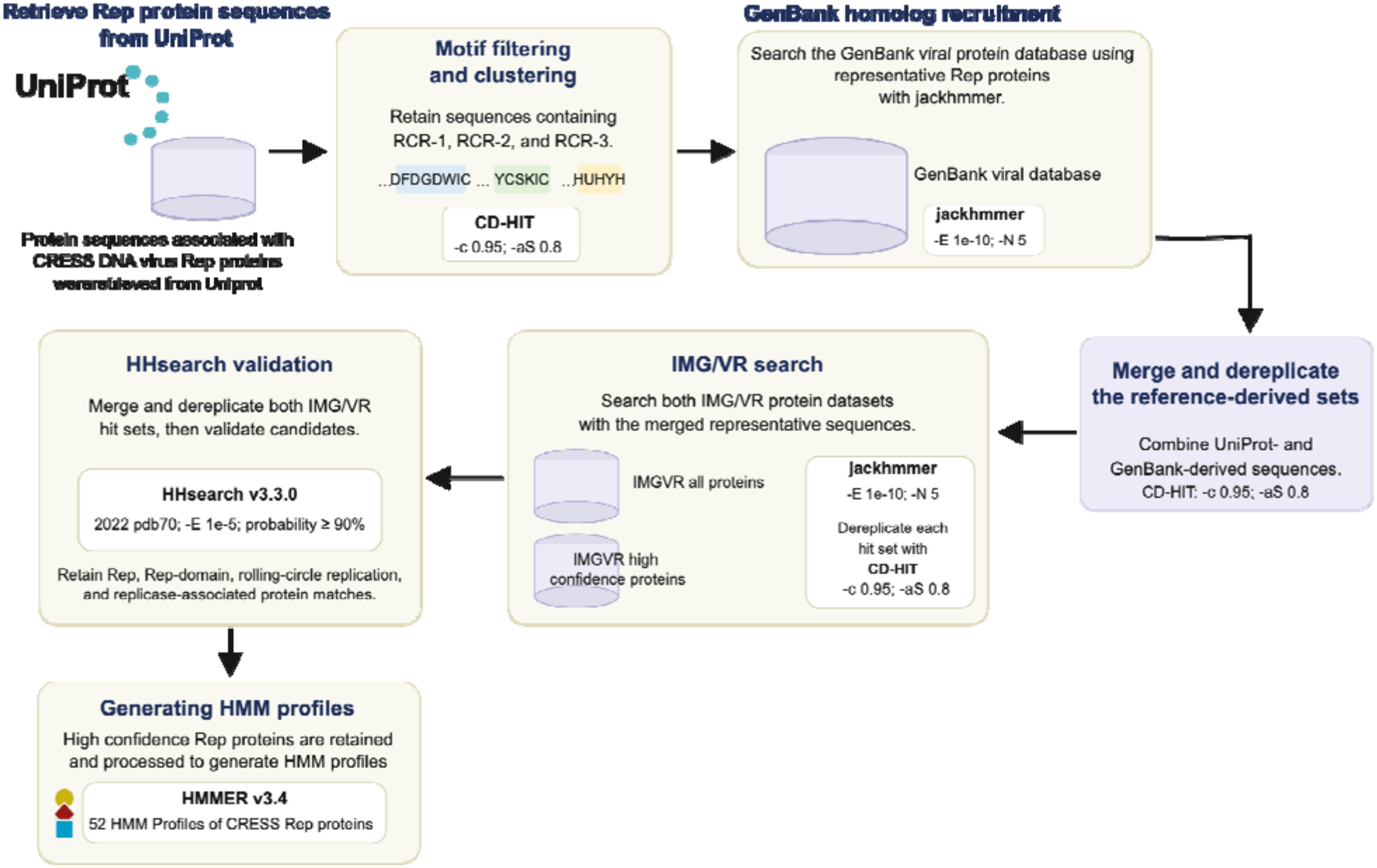
Workflow for the identification of CRESS DNA virus replication-associated proteins (Rep). The workflow illustrates the retrieval, dereplication, and clustering of Rep sequences from UniProt, GenBank, and IMG/VR datasets, followed by validation using HHsearch, and the construction of 52 hidden Markov model (HMM) profiles.

#### Acquisition and processing of Capsid protein sequences

The 11,892 viral sequences belonging to Cressdnaviricota retrieved from UniProt, which were determined to possess complete sequences of replication-associated proteins (Rep) and conserved rolling-circle replication motifs RCR-1, RCR-2, and RCR-3, were further used to retrieve sequences annotated as major capsid proteins. From these, a total of 10391 complete capsid sequences were retrieved and dereplicated into 2,848 representative sequences with cd-hit v.4.8.1 (-c 0.95 and -aS 0.8) and subsequently validated as viral capsid proteins with HHsearch v.3.3.0 against the 2023 UniRef30 database (-E 1e-5).

#### Generating HMM profiles for replication-associated proteins

The retained set of 25,416 high-confidence Rep proteins was combined with 7,582 reference sequences obtained from UniProt and GenBank and compared using reciprocal BLAST v.2.13.0+ (1e-10). Sequences were subsequently grouped into clusters using the Markov Cluster Algorithm v.14-137 (MCL) with an inflation parameter of 2. Sequences within each cluster were aligned using MAFFT L-INS-i v7.505 and trimmed with TrimAl v.1.4.rev15 (gappyout option). The resulting trimmed alignments were then validated as Rep-associated through HHsearch v3.3.0 against the PDB70 (2022) database. Validated clusters were subsequently used as input for HMMER v3.4 to generate a total of 52 HMM profiles.

#### Generating HMM profiles for major Capsid proteins

The dereplicated set of 2,848 major capsid protein sequences were compared against each other using reciprocal BLAST v.2.13.0+ (1e-10) and further clustered using MCL (Enright et al., 2002) with the inflation parameter set to 2. Sequences within each these clusters were also aligned using MAFFT L-INS-i v7.505, trimmed with TrimAl v.1.4.rev15 (gappyout option), and alignments further validated as viral capsid-associated through HHsearch v3.3.0 against the UniRef30 (2023) database. Validated alignments were subsequently used as input for HMMER v3.4 to generate a total of 32 HMM profiles.

#### Screening for Endogenous Viral Elements

We screened 1,599 fungal genomes from MycoCosm and 194 algal genomes from PhycoCosm (both accessed February 2025) for genome-associated Rep-like loci using the 52 HMM profiles and hmmscan v3.4 (-T 50). Rep proteins predicted in these genomes were validated as Rep-associated using HHsearch against the PDB70 database. Contigs predicted to encode these genes were additionally screened for the co-localization of genes associated with major capsid proteins. The screening for capsid proteins was conducted using a set of 36 HMM profiles with hmmscan v3.4 (-T 50) and HHblits v3.3.0 against the UniRef30 (2023) database.

#### Phylogenetic Analysis of Rep Proteins

For phylogenetic analysis, we combined 8,286 high-confidence Rep proteins (a set retained from IMGVR_all_proteins_high_confidence.faa), 1,876 ICTV MSL40.v1 Cressdnaviricota representatives, and 1,025 genome-associated Rep-like sequences. These were aligned using MAFFT L-INS-i v7.505 and subsequently trimmed with TrimAl (gappyout parameter). The trimmed alignment was then used to reconstruct a maximum-likelihood phylogenetic tree in IQ-TREE v3.0.1 (-m MFP -bb 1000 -alrt 1000 parameters). Based on the resulting phylogenetic tree, we clustered Rep sequences using PDMclust, a phylogeny-based pairwise distance matrix (PDM) approach (https://github.com/NeLLi-team/pdmclust#). This yielded 6,391 total clusters, of which 4,650 were singletons. After excluding singletons, we selected one representative from each of the 1,741 clusters inferred using PDM. We realigned the 1,741 representatives with MAFFT L-INS-i v7.505, trimmed with TrimAl (gappyout setting), and inferred the final maximum-likelihood tree in IQ-TREE v3.0.1 (-m MFP -bb 1000 -0alrt 1000). The resulting tree file was visualized using iTOL, and clades were annotated based on family-level classifications obtained from the ICTV taxonomy database.

## Results and Discussion

### Recovery of environmental Rep protein diversity

Following iterative sequence searches, a diverse and non-redundant set of 25,416 Rep proteins was recovered from IMG/VR data. The large number of Rep proteins identified highlights the extensive and still poorly characterized diversity of CRESS DNA viruses across ecologically diverse metagenomic samples. Similar observations have been reported in other metagenomic studies, where environmental sequencing uncovered a variety of unclassified and highly divergent CRESS DNA viruses across host-associated and environmental samples (Bezuidt and Makhalanyane, 2024, Trubl et al., 2019, van der Loos et al., 2023). For example, Liu et al. (2020) identified eight novel CRESS DNA virus genomes from forest musk deer (Liu et al., 2020), while Sommers et al. (2019) identified seven viral sequences representing previously unclassified CRESS viruses from Antarctic cryoconite (Sommers et al., 2019). Notably, Tisza et al. (2020) conducted one of the largest systematic surveys of circular DNA viruses in animal-associated samples, recovering 2,514 complete genomes, of which 868 were associated with unclassified CRESS DNA viruses (Tisza et al., 2020). A study by Trubl et al. (2025) also identified 574 viral operational taxonomic units (vOTUs) associated with CRESS DNA viruses from permafrost, and reported that 99% of these were unclassified CRESS viruses (Trubl et al., 2025). Together, these findings support the growing view that environmental and host-associated viromes represent a major source of unexplored ssDNA viral diversity.

### Identification of endogenous viral elements in fungal and algal genomes

Screening of 1,599 fungal and 194 algal genomes identified 1,025 CRESS DNA virus-associated Rep-like loci. Given that Rep-like sequences have previously been used as evidence of CRESS DNA virus-derived integrations in eukaryotic genomes, these sequences were considered candidate endogenous viral elements, hereafter referred to as EVEs (Katzourakis and Gifford, 2010, Liu et al., 2011, Zhao et al., 2021). Of the total predicted Rep-like sequences, 751 were predicted in 132 fungal genomes and 274 in 24 algal genomes (**Figure S1A and Supplementary Table 1**). The number of Rep-like loci per genome ranged from 1 to 100. Among fungal genomes, the largest numbers of EVEs were identified in *Gigaspora margarita* BEG34, *Zychaea mexicana* RSA 1403 v1.0, and *Gigaspora rosea* v1.0, with 100, 86, and 62 Rep-like loci, respectively (**Figure 2A**). In algal genomes, the highest numbers of EVEs were identified in *Seminavis robusta* D6, *Polarella glacialis* CCMP1383, and *Polarella glacialis* CCMP2088, which contained 72, 56, and 32 Rep-like sequences, respectively (**Figure 2B**). Among these, several EVE loci were co-located with predicted major capsid genes, consistent with the retention of multiple viral genes within some host chromosomes (**Figure 2C and D**). In total, 100 capsid genes were identified within EVEs across 19 fungal genomes, compared with 28 capsid genes detected within EVEs across 10 algal genomes. Similar endogenous viral sequences have previously been identified in animal and plant genomes, where they have provided evidence of long-term virus–host interactions and viral evolution (Ter Horst et al., 2019, Chu et al., 2014, Katzourakis and Gifford, 2010).

**Figure 2:**
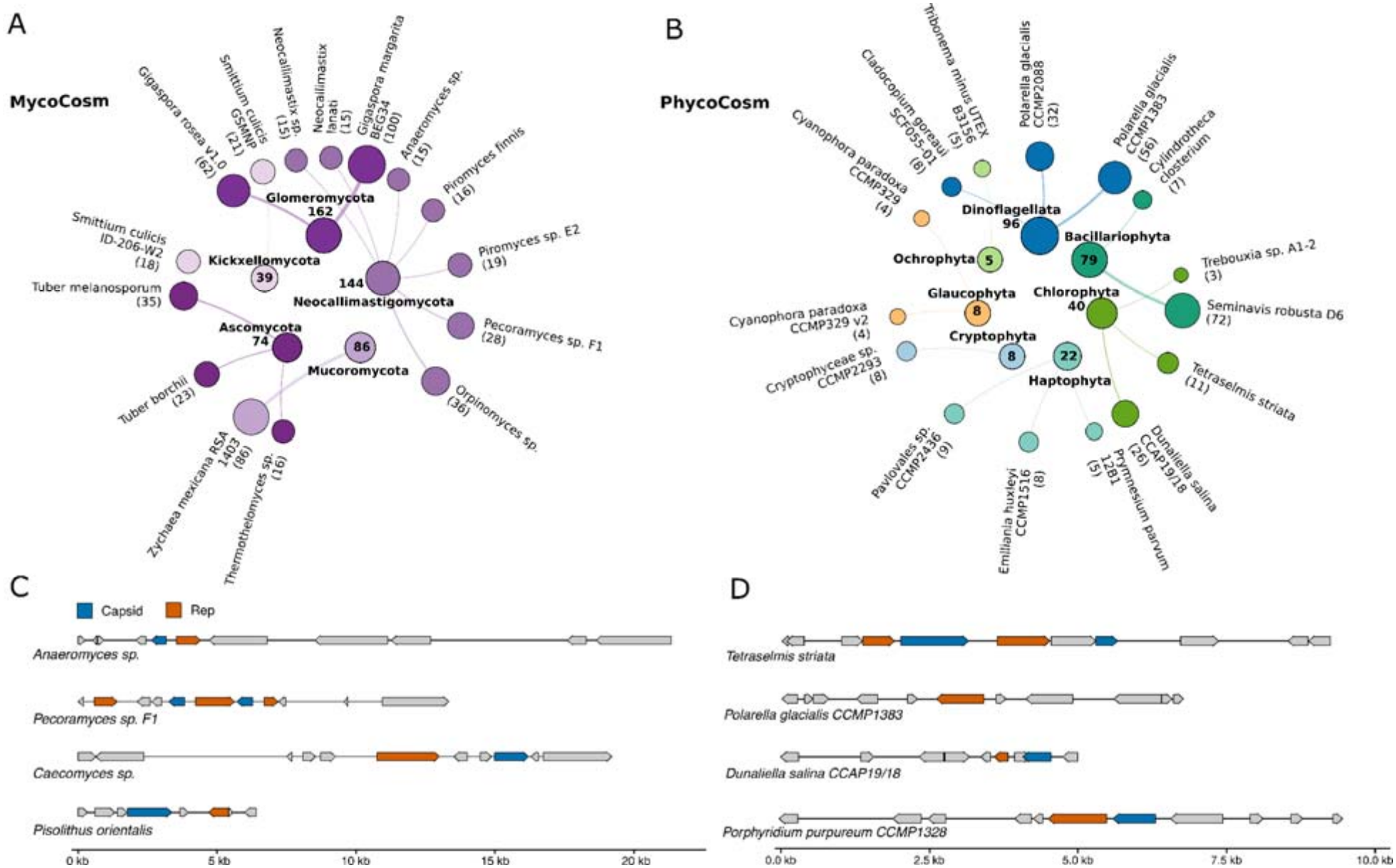
Distribution of endogenous viral elements (EVEs) in fungal and algal genomes. Screening of MycoCosm and PhycoCosm genomes identified 751 EVEs across fungal genomes and 274 EVEs across algal genomes, respectively. The figure illustrates the top 15 (**A**) fungal and (**B**) algal genomes containing the highest numbers of predicted EVEs. The maps illustrate the genomic organization of capsid and CRESS DNA virus replication-associated protein (Rep) genes and their neighbouring host genes predicted in (**C**) fungal and (**D**) algal genomes. Arrows indicate gene orientation and relative genomic positions, with Rep genes highlighted in orange, capsid genes in blue, and neighbouring host genes in grey. Genomic coordinates are shown in kilobases (kb).

Compared with animals and plants, fungal and algal genomes remain comparatively underexplored for viral endogenization events. Nevertheless, several reports have suggested that fungal-associated viruses and EVE may be more widespread than previously recognized (Myers et al., 2026, Kerr et al., 2018, Liu et al., 2016, Zhao et al., 2023, Clemons et al., 2024). Yet, reports of endogenous ssDNA virus-derived elements in fungal genomes remain sparse, suggesting either limited integration frequency or insufficient genomic exploration of fungal viromes.

Algal systems appear to show a different pattern, with studies reporting widespread viral endogenization events in marine and freshwater algae (Duchene et al., 2026a, Duchene et al., 2026b, Jivaji et al., 2025). Although many of these studies focused on double-stranded DNA viruses of the phyla Nucleocytoviricota and Mirusviricota, as well as Preplasmaviricota rather than CRESS DNA viruses, they collectively demonstrate that algal genomes act as important reservoirs of endogenous viral diversity (Chase et al., 2024, Gyaltshen et al., 2023, Nelson et al., 2021). The detection of CRESS DNA virus-derived EVEs in fungal and algal genomes in this study therefore broadens the potential host associations of Cressdnaviricota and suggests that these organisms may represent overlooked hosts or historical reservoirs of ssDNA viruses.

### Phylogenetic diversity of environmental and endogenous Rep proteins

A maximum-likelihood phylogenetic tree using 1,736 representative Rep proteins from IMG/VR, EVEs, and ICTV MSL40.v1 revealed extensive diversity across CRESS DNA viruses (**Figure 3**). To assess the diversity and taxonomic placement of newly identified Rep sequences, phylogenetic clades were evaluated using ultrafast bootstrap (UFBoot) support values ≥95 and SH-aLRT support values ≥80 to facilitate taxonomic classification. We provisionally associated non-ICTV Rep sequences with an ICTV family when they occurred in well-supported clades containing ICTV reference sequences. In addition to these classified clades, well-supported branches lacking ICTV representatives, containing five or more sequences, supported by UFBoot ≥95 and SH-aLRT ≥80, were considered to represent divergent evolutionary lineages (candidate novel families) not currently represented in ICTV classifications. Of the 1,409 non-ICTV tips, 603 (43%) were provisionally associated with an ICTV family and 806 (57%) were not classified with ICTV families (Figure S1B). Of the latter, 479 tips (59.4%) formed 52 unclassified monophyletic clades containing ≥5 viruses (supported by UFBoot ≥95 and SH-aLRT ≥80) falling outside the currently recognized ICTV taxonomic framework. Many of these sequences came from freshwater (*n=308*) and marine *(n=68*) samples (Figure 3). The 52 unclassified monophyletic clades suggest that these viruses could potentially expand the diversity of recognized CRESS families, with some highlighted in the tree (**Figure 3**) as new CRESS lineages (NCLs) 1–16. The tree also illustrates polyphyletic groupings within established families, such as *Adamaviridae, Circoviridae*, and *Draupnirviridae*, which could potentially be further resolved into distinct CRESS families. Several other distinct clades that may represent potentially new CRESS families are highlighted in the tree as NCL1, NCL3, NCL5, and NCL14–16.

**Figure 3:**
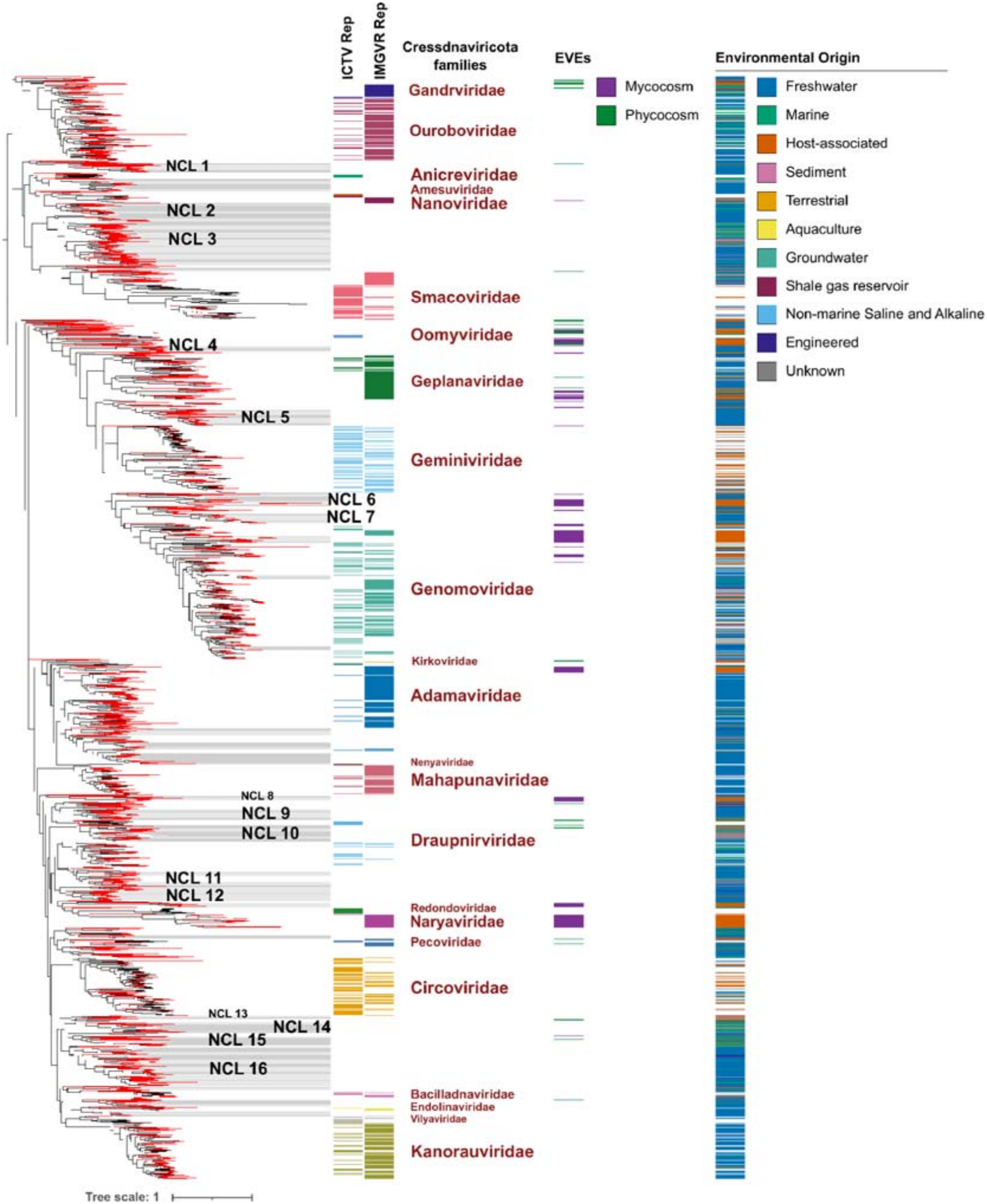
Maximum-likelihood phylogenetic tree of CRESS DNA virus replication-associated proteins (Rep). The tree was inferred from representative Rep protein sequences retrieved from IMG/VR, ICTV reference viruses, and endogenous viral elements (EVEs) identified in fungal and algal genomes. Red branches represent Non-ICTV sequences (IMG/VR and EVEs), whereas black branches represent sequences retrieved from ICTV. Non-ICTV sequences were provisionally associated with established ICTV families, while supported unclassified monophyletic clades, shaded in grey, were designated as potentially novel CRESS lineages (NCLs).

EVE-derived representative Rep proteins (*n=153*) were distributed across multiple regions of the phylogenetic tree rather than clustering within a single lineage, suggesting that endogenous sequences reflect broad evolutionary diversity across Cressdnaviricota. Of these, 97 (63.4%) sequences were unclassified, whereas 56 (36.6%) grouped in monophyletic clades of known ICTV families (Figure 3). MycoCosm EVE-derived Rep proteins (50 of 118, 42.4%) grouped with *Naryaviridae* (*n=20*), *Geplanaviridae* (*n=11*), *Adamaviridae* (*n=9*), *Genomoviridae* (*n=9*), and *Nanoviridae* (*n=1*). This grouping highlights the complexity and diversity of CRESS viruses associated with fungi, contrasting with studies that linked *Naryaviridae* to parasites of the genus *Entamoeba* (Krupovic and Varsani, 2022, Kinsella et al., 2020). This underlines that eukaryotic genomes may preserve evolutionary lineages absent from current viral reference databases.

Together, these findings demonstrate that integrating environmental metagenomic datasets with endogenous viral element analyses broadens our understanding of the diversity and evolutionary history of CRESS DNA viruses. While environmental viromes capture extant viral diversity, EVEs preserve evidence of historical virus–host interactions, collectively contributing to a broader understanding of the ecological distribution, host associations, and long-term evolution of ssDNA viruses.

## Acknowledgements

The work conducted by the U.S. Department of Energy Joint Genome Institute (https://ror.org/04xm1d337), a DOE Office of Science User Facility, is supported by the Office of Science of the U.S. Department of Energy operated under Contract No. DE-AC02-05CH11231.

## Data availability

Files generated in this study are available from: **10.5281/zenodo.24107104**. These include the HMM profiles generated to screen for Rep-encoding proteins and major capsid proteins, a spreadsheet with IDs of MycoCosm and PhycoCosm genomes screened for EVEs, FASTA files containing Rep protein sequences representing EVEs predicted in MycoCosm and PhycoCosm genomes, and files indicating the number of EVEs identified in fungal and algal hosts, together with the taxonomic lineages of the respective host genomes.

## Author contributions

O.K.B. and F.S. designed the study, coordinated all analyses, and provided overall project oversight. Both authors analyzed the data and contributed to writing the manuscript.

## Figure legends

**Supplementary Figure 1:**
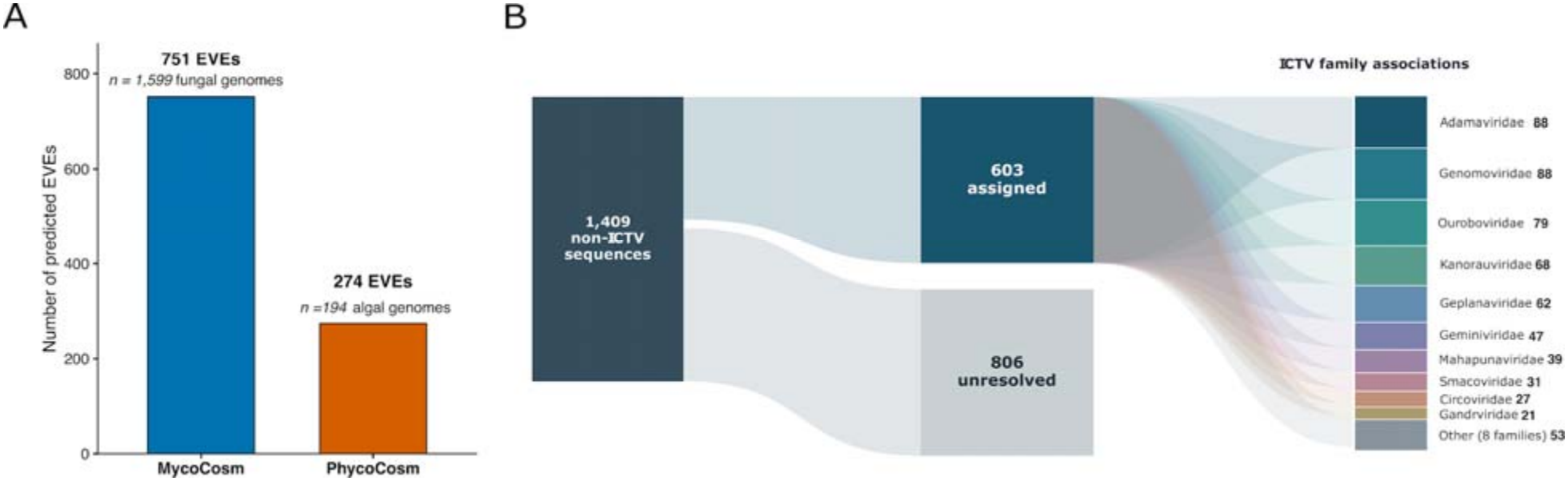
Distribution of EVEs and family classifications of Rep-encoding proteins. (**A**) Number of EVEs identified in fungal and algal genomes from the MycoCosm and PhycoCosm databases, respectively. (**B**) Flow diagram illustrating the provisional association of non-ICTV Rep protein sequences with established ICTV families.

